# Microbiota-derived indole-3-propionic acid regulates glucose homeostasis via remodeling of hepatic mitochondrial metabolism

**DOI:** 10.64898/2026.05.11.724210

**Authors:** Or Maalumi, Ziv Ben Moshe, Or Blank, Rachel Barkan-Michaeli, Avihu Yona, Kfir Sharabi

## Abstract

The gut microbiota produces metabolites that circulate to host tissues and are increasingly linked to metabolic health, yet the mechanisms by which individual microbial products regulate liver glucose metabolism remain poorly defined. Here, we identify the tryptophan-derived microbial metabolite indole-3-propionic acid (IPA) as a direct modulator of hepatic glucose production. In primary hepatocytes, a focused screen of indole metabolites revealed that several indole-containing compounds suppress glucagon-stimulated glucose output, with IPA emerging as a physiologically relevant candidate. IPA selectively reduced glucose production from mitochondrial-dependent gluconeogenic substrates while largely preserving glycerol-supported glucose production, suggesting that it does not simply shut down gluconeogenesis but instead alters how hepatocytes use metabolic fuels. Mechanistic analyses showed that IPA redirects lactate-derived carbon away from glucose production and reshapes mitochondrial metabolism, including redox balance, ATP availability, and urea cycle-linked metabolic activity. These effects occurred without detectable disruption of proximal insulin or glucagon signaling, supporting a model in which IPA acts primarily through intracellular metabolic remodeling. In mice, endogenous IPA levels varied with nutritional state, and short-term IPA administration improved fasting glycemia and glucose handling in Western diet-fed animals. Finally, microbiome-depleted mice colonized with IPA-producing *Clostridium sporogenes* displayed increased circulating IPA and improved glucose tolerance compared with mice colonized with an IPA-deficient mutant *C. Sporogenes* strain. Together, these findings identify IPA as a microbial metabolite that directly connects gut tryptophan metabolism to hepatic mitochondrial function and systemic glucose regulation, highlighting a mechanistic gut-liver pathway with potential therapeutic relevance to metabolic disease.

## Introduction

Maintenance of fasting blood glucose levels relies predominantly on hepatic glucose production (HGP), a process tightly regulated by the opposing actions of insulin and glucagon^1–3^. In insulin-resistant states, including metabolic dysfunction-associated steatotic liver disease (MASLD) and type 2 diabetes, suppression of HGP is impaired, leading to excessive hepatic glucose output and fasting hyperglycemia^4,5^. While transcriptional induction of gluconeogenic enzymes contributes to this dysregulation, accumulating evidence indicates that acute, post-translational, and metabolic control mechanisms play a major role in determining gluconeogenic flux^6,7^. Glucagon is a dominant driver of HGP during fasting and in insulin-resistant conditions, acting through cAMP-dependent signaling pathways to stimulate gluconeogenesis and hepatic energy mobilization. Importantly, glucagon-stimulated glucose production depends not only on canonical signaling cascades but also on the metabolic capacity of hepatocytes to support sustained carbon flux through gluconeogenic pathways^8–10^. How this metabolic capacity is modulated in physiological and pathophysiological contexts remains incompletely understood.

The gut microbiota has emerged as a critical regulator of host metabolic homeostasis through the production of bioactive metabolites that enter the systemic circulation^11,12^. These metabolites, derived from dietary components and microbial metabolic activity, can influence glucose and lipid metabolism, inflammation, and energy balance across multiple organs^13^. Because the liver is directly exposed to gut-derived metabolites via the portal vein, hepatocytes are uniquely positioned to integrate microbial signals into metabolic responses^14^. Despite growing recognition of microbiota-host crosstalk, the mechanisms by which individual microbial metabolites influence hepatic metabolic pathways remain poorly defined. In particular, whether these metabolites exert their effects primarily through receptor-mediated signaling, transcriptional reprogramming, or direct modulation of intracellular metabolic processes is often unclear. Dissecting these mechanisms is essential for understanding how gut microbial activity contributes to metabolic health and disease.

Indole metabolites constitute a chemically related group of microbiota-derived compounds produced from dietary tryptophan and present in the systemic circulation^15,16^. Several indoles reach the liver via the portal vein, positioning hepatocytes as direct targets of microbial metabolic activity^15^. Within this class, indole-3-propionic acid (IPA) has emerged as a prominent metabolite due to its abundance, microbial origin, and consistent association with favorable metabolic phenotypes. In experimental models, IPA improves glucose metabolism in rats^17^, protects intestinal barrier function through PXR- and TLR4-linked mechanisms^18^, attenuates diet-induced steatohepatitis in rats by reducing gut dysbiosis and endotoxin leakage^19^, and protects against heart failure with preserved ejection fraction, in part through effects on NAD metabolism^20^. More recent studies further support the idea that IPA can act through metabolic remodeling: IPA promotes mitochondrial respiration in CD4+ T cells to confer protection against intestinal inflammation^21^, and microbiota-derived IPA promotes mucosal healing in colitis through intestinal HMGCS2-dependent ketogenesis^22^. Together, these studies position IPA as a bioactive microbial metabolite capable of influencing host metabolism, mitochondrial function, inflammatory tone, and tissue homeostasis across multiple physiological contexts.

Despite this growing body of evidence, the direct hepatic actions of IPA remain insufficiently defined. Prior studies in metabolic disease models have largely emphasized systemic, intestinal, inflammatory, or microbiome-mediated mechanisms, whereas it remains unclear whether IPA can act directly on hepatocytes to regulate glucose production. This distinction is particularly important because hepatic glucose output is acutely controlled by hormone-driven metabolic flux, not only by longer-term transcriptional or inflammatory remodeling. Although IPA has been implicated in signaling through host receptors such as the aryl hydrocarbon receptor and pregnane X receptor^18,23,24^, and in the modulation of mitochondrial or NAD-linked metabolism in other cell types^20,21^, whether these mechanisms converge on hepatocyte gluconeogenic control during glucagon stimulation has not been established. Specifically, it remains unknown whether IPA influences hepatic glucose output through altered hormone signaling, changes in substrate handling, or direct remodeling of intracellular metabolic state.

Hepatic glucose production is ultimately constrained by mitochondrial metabolic capacity, which integrates hormonal cues with intracellular redox balance, energy state, and carbon flux through central metabolic pathways. During glucagon stimulation, efficient mitochondrial function is required to sustain gluconeogenic flux even in the presence of intact upstream signaling. This study was motivated by previous work identifying SR18292 as a small-molecule suppressor of glucagon-induced hepatic glucose production^25^, followed by structure-activity analyses demonstrating that its indole-containing chemical scaffold is required for this biological activity^26^. These findings raised the possibility that structurally related indole-containing metabolites, including microbiota-derived indoles, may also influence hepatic glucose production. We therefore performed a targeted screen of selected indole metabolites and identified IPA as a biologically relevant suppressor of glucagon-stimulated glucose production. We then used mechanistic analyses in primary hepatocytes, including metabolic profiling, isotope tracing, and measurements of cellular redox and energy status, to define how IPA alters hepatocyte metabolic function under glucagon-stimulated conditions. These in vitro studies were complemented by acute in vivo IPA treatment in mice to assess the physiological relevance of our findings for whole-body glucose homeostasis. Together, this integrated approach was designed to determine whether IPA functions as a microbiota-derived effector linking gut microbial metabolism to hepatic glucose regulation.

## Results

### Indole derivatives suppress hepatic glucose production in a substrate-selective manner

Guided by prior structure-activity relationship studies demonstrating that the indole moiety of SR-18292 is required for its ability to suppress hepatic glucose production^25,26^, we asked whether structurally related indole-containing metabolites can directly influence hepatocyte glucose output. To test this, we performed a focused screen of selected indole derivatives (Fig. 1A) and measured glucagon-stimulated glucose production in primary hepatocytes using pyruvate/lactate or glycerol as gluconeogenic substrates. Several indole derivatives significantly reduced glucose production from pyruvate/lactate, whereas glucose production from glycerol was less consistently affected across the compound panel (Fig. 1B). These findings indicate that indole derivatives do not act as broad inhibitors of gluconeogenic capacity, but instead suppress glucose production in a substrate-selective manner.

**Figure 1.**
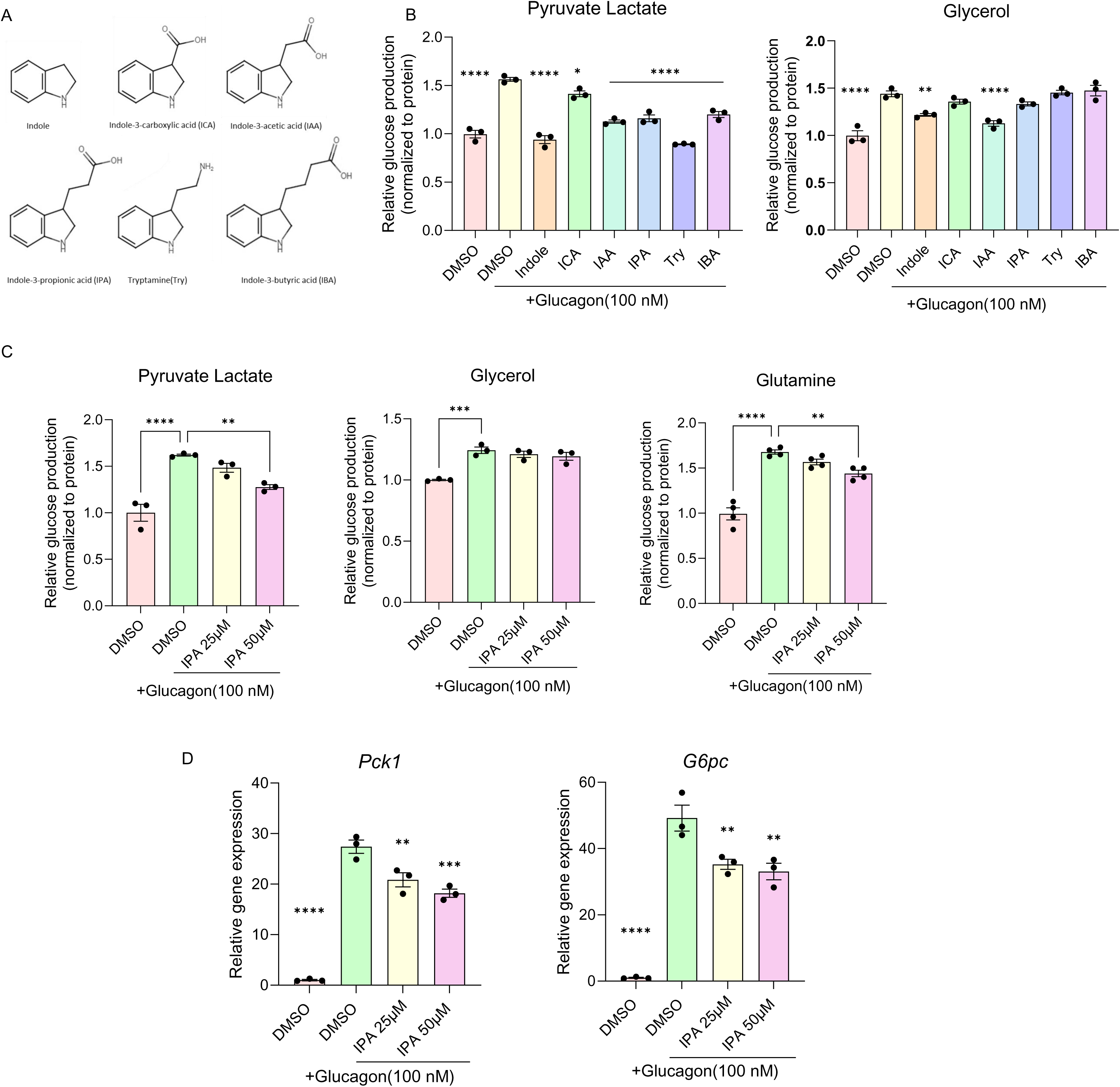
Indole-derived metabolites suppress hepatic glucose production. **(A)** Chemical structure of the indole derivatives that were tested. **(B)** Hepatic glucose production (HGP) in primary hepatocytes treated with indole-derived metabolites using pyruvate/lactate or glycerol as substrates. For statistics, the comparison was made to DMSO + glucagon conditions. **(C)** HGP in primary hepatocytes treated with increasing concentrations of indole-3-propionic acid (IPA) in the presence of pyruvate/lactate, glycerol, or glutamine. **(D)** Expression of gluconeogenic genes in primary hepatocytes following IPA treatment. *, p<0.05; **, p<0.01; ***, p<0.001; ****, p<0.0001.

Among the compounds tested, we focused on indole-3-propionic acid (IPA), a microbiota-derived indole metabolite associated with favorable metabolic phenotypes in humans^27–29^. Dose-response experiments confirmed that IPA suppresses glucagon-stimulated glucose production from pyruvate/lactate, while glucose production from glycerol was largely preserved over the same concentration range (Fig. 1C). Notably, IPA also reduced glucose production from glutamine, particularly at the higher concentration tested (Fig. 1C). Thus, IPA preferentially suppresses glucose production from substrates that require mitochondrial metabolism or anaplerotic processing before entering the gluconeogenic pathway, while sparing glycerol-supported glucose production. Consistent with the reduction in glucose output, IPA also dose-dependently attenuated glucagon-induced expression of the canonical gluconeogenic genes *Pck1* and *G6pc* (Fig. 1D). These data indicate that IPA can suppress both functional glucose production and selected components of the glucagon-induced gluconeogenic transcriptional program. However, because glycerol-supported glucose production was largely preserved despite reduced *Pck1* and *G6pc* expression, the functional effect of IPA cannot be explained solely by broad transcriptional repression of gluconeogenesis. We next examined whether this transcriptional effect extended more broadly across the indole derivative panel. Glucagon robustly induced *G6pc* and *Pck1* expression, and several indole derivatives attenuated this response, with the most consistent suppression observed for *Pck1* (Fig. S1A). In contrast, effects on *G6pc* were more variable, and *Ppargc1a* expression was not uniformly reduced across the compound panel. These data indicate that indole derivatives can partially suppress selected components of the glucagon-responsive gluconeogenic gene program, but do not cause a broad shutdown of glucagon-induced transcription.

Because IPA was selected for further mechanistic analysis, we next asked whether its effect could be explained by altered proximal hormone signaling. IPA did not reduce insulin-stimulated AKT phosphorylation, indicating that canonical insulin signaling remained intact (Fig. S1B). Similarly, IPA did not blunt glucagon-induced CREB phosphorylation, suggesting that suppression of glucose production is not due to inhibition of proximal glucagon-CREB signaling (Fig. S1C). Together, these findings support the idea that IPA acts downstream of, or parallel to, canonical hormone receptor signaling.

IPA has also been reported to signal through host xenobiotic receptors, including the aryl hydrocarbon receptor (AhR) and the pregnane X receptor (PXR/*Nr1i2*)^18,23,24^, which mediate some of its biological effects in other tissues and experimental systems^18,23^. We therefore tested whether these pathways are required for IPA-mediated suppression of glucose production. Pharmacological inhibition of AhR with CH223191 did not prevent IPA from reducing glucagon-stimulated glucose production (Fig. S2A). Similarly, siRNA-mediated knockdown of *Nr1i2*, which encodes PXR, did not rescue glucose output in IPA-treated hepatocytes (Fig. S2B). IPA also attenuated glucagon-induced *Pck1* and *G6pc* expression in control conditions, although the effect on these transcripts was not uniformly preserved across receptor-perturbation conditions (Fig. S2B, D). Thus, while the transcriptional response may vary depending on receptor pathway perturbation, the acute suppression of glucagon-stimulated glucose production by IPA does not require AhR or PXR/*Nr1i2*.

Together, these data identify IPA as a microbiota-derived indole metabolite that suppresses glucagon-stimulated hepatic glucose production in a substrate-selective manner. The preferential effect on pyruvate/lactate-driven glucose output, together with preserved glycerol-supported glucose production and largely intact proximal hormone signaling, suggested that IPA does not simply inhibit gluconeogenesis globally. Instead, these findings pointed to a more specific effect on the handling of lactate-derived carbon during glucagon-stimulated glucose production.

### IPA remodels lactate-derived carbon flux at the TCA-urea cycle interface

Having established that IPA suppresses glucagon-stimulated glucose production from pyruvate/lactate, we next asked whether this phenotype reflects altered handling of lactate-derived carbon. To address this, we performed metabolic tracing using uniformly labeled [U-^13^C]-lactate in primary hepatocytes under basal and glucagon-stimulated conditions. Consistent with the reduction in glucose output observed functionally, IPA significantly reduced the abundance of ^13^C□-labeled glucose-6-phosphate (G6P), particularly under glucagon-stimulated conditions (Fig. 2A,B). This indicates that IPA limits the incorporation of lactate-derived carbon into the gluconeogenic pathway.

**Figure 2.**
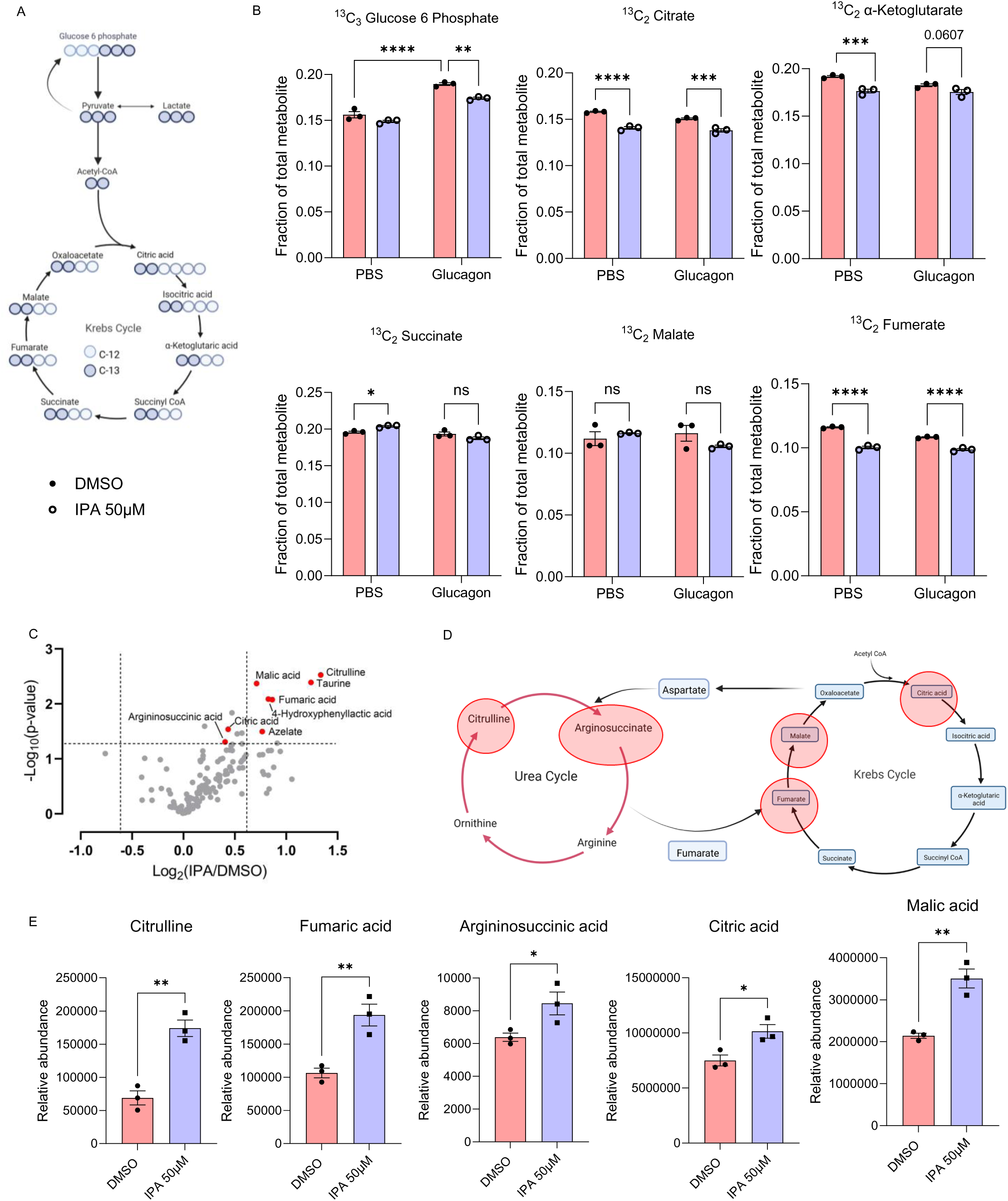
IPA rewires hepatic carbon metabolism. **(A)** Schematic of ^13^C□-lactate tracing into gluconeogenic and TCA cycle intermediates. **(B)** Fractional enrichment of ^13^C-labeled metabolites in primary hepatocytes. **(C-D)** Untargeted metabolomic analysis identified several metabolites at the TCA and urea cycles intersection, that are altered in response to acute IPA treatment. **(E)** Metabolite changes at the intersection of the TCA and urea cycles. *, p<0.05; **, p<0.01; ***, p<0.001; ****, p<0.0001., ns, not significant.

We then examined whether IPA also alters the propagation of lactate-derived carbon through mitochondrial metabolic intermediates. Across several TCA cycle metabolites, including citrate, α-ketoglutarate, malate, and fumarate, IPA reduced ^13^C□ enrichment from lactate in both basal and glucagon-stimulated states (Fig. 2B). This coordinated reduction across multiple intermediates argues against inhibition at a single enzymatic step and instead suggests that IPA broadly alters the entry or retention of lactate-derived carbon within mitochondrial carbon metabolism. Notably, IPA reduced labeling of malate and fumarate, intermediates positioned at the interface between the TCA cycle, gluconeogenesis, and nitrogen metabolism, suggesting that IPA affects metabolic nodes directly relevant to lactate-supported glucose production.

To determine whether IPA also induces broader metabolic remodeling independent of acute hormonal stimulation, we performed untargeted metabolomics in primary hepatocytes treated with IPA for 3 h under basal conditions. Volcano plot analysis revealed a focused set of metabolites increased by IPA rather than a global shift across the metabolome (Fig. 2C). Pathway mapping showed that the most responsive metabolites clustered at the intersection of mitochondrial carbon metabolism and nitrogen handling (Fig. 2D). IPA increased the abundance of urea cycle-associated intermediates, including citrulline and argininosuccinic acid, together with connected TCA cycle intermediates, including fumaric acid, malic acid, and citric acid (Fig. 2E). Citrulline showed the largest increase, while fumarate and malate were also prominently elevated. Importantly, these steady-state metabolite changes were observed in the absence of glucagon stimulation, indicating that IPA induces an early reorganization of hepatocyte metabolic state rather than simply blocking glucagon signaling. When considered together with the isotope-tracing data, the accumulation of selected TCA-urea cycle intermediates despite reduced lactate-derived labeling suggests that IPA alters the source or turnover of these metabolite pools, rather than uniformly suppressing mitochondrial metabolism. Thus, IPA limits the contribution of lactate-derived carbon to gluconeogenic and mitochondrial intermediates while selectively remodeling metabolite pools at the TCA-urea cycle interface.

Together, these findings provide a metabolic explanation for the substrate-selective suppression of glucose production observed in Figure 1. IPA does not appear to broadly inhibit gluconeogenesis, but rather alters the metabolic routing of lactate-derived carbon through pathways that support glucagon-stimulated glucose production.

### IPA reshapes mitochondrial redox state and urea cycle-linked metabolism during glucagon stimulation

Consistent with the reduced incorporation of lactate-derived carbon into gluconeogenic and mitochondrial intermediates observed in the ^13^C-lactate tracing experiments above, we next asked whether IPA alters hepatocyte redox state and energy balance during glucagon stimulation. IPA treatment increased the cellular NAD□/NADH redox balance relative to vehicle control (Fig. 3A). Glucagon stimulation similarly elevated the NAD□/NADH redox balance, consistent with activation of oxidative metabolism during gluconeogenic stimulation. However, the combination of IPA and glucagon did not produce a further increase and instead shifted the ratio toward baseline levels, indicating that IPA modifies the redox response to glucagon rather than simply amplifying mitochondrial oxidation.

**Figure 3.**
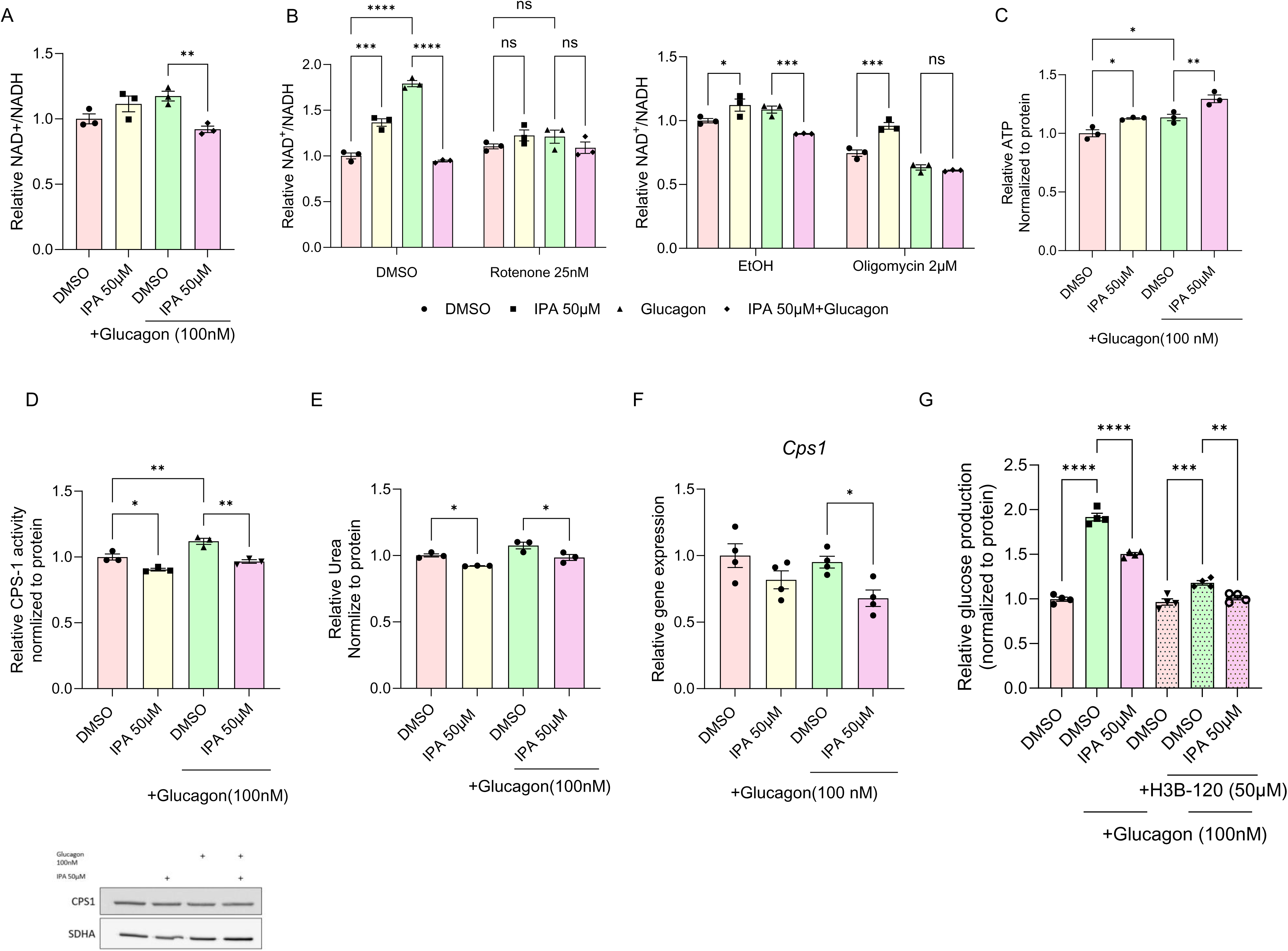
IPA regulates hepatic redox and ureagenesis. **(A)** NAD□/NADH ratio under IPA treatment glucagon stimulation. **(B)** NAD□/NADH ratio following inhibition of Complex I (rotenone) or Complex V (oligomycin) of the mitochondrial electron transport chain. **(C)** Intracellular ATP levels. **(D)** CPS1 enzymatic activity and western blot analysis of protein abundance. **(E)** Urea secretion. **(F)** *Cps1* gene expression. **(G)** Hepatic glucose production in the presence of H3B-120 (CPS1 inhibitor) and IPA. *, p<0.05; **, p<0.01; ***, p<0.001; ****, p<0.0001; ns, not significant.

To determine whether these redox effects depend on mitochondrial respiration, we repeated these measurements in the presence of respiratory inhibitors. Inhibition of Complex I with rotenone abolished the IPA-induced increase in NAD□/NADH redox balance and also prevented the glucagon-induced increase observed under control conditions (Fig. 3B). Under rotenone treatment, neither glucagon nor glucagon + IPA increased the NAD□/NADH ratio relative to basal conditions, indicating that both the IPA-dependent redox shift and the glucagon-associated redox response require Complex I-dependent mitochondrial NADH oxidation. This suggests that IPA does not act by independently raising NAD□/NADH downstream of respiratory blockade, but rather modifies redox balance through Complex I-linked respiration. In contrast, inhibition of ATP synthase with oligomycin did not prevent the IPA-induced increase in NAD□/NADH redox balance under basal conditions, but it abolished the glucagon-associated increase in NAD□/NADH and eliminated the effect of IPA under glucagon stimulation (Fig. 3B). Thus, the basal redox effect of IPA can still occur when ATP synthase is inhibited, whereas the glucagon-dependent redox response requires intact oxidative phosphorylation. In parallel, IPA increased cellular ATP levels under both basal and glucagon-stimulated conditions (Fig. 3C), indicating that IPA does not broadly impair mitochondrial energy production. Because ATP increased in both conditions whereas the NAD□/NADH response differed between basal and glucagon-stimulated states, ATP levels do not simply mirror the redox ratio. Rather, these data suggest that IPA alters mitochondrial energy handling together with Complex I-dependent redox balance, with glucagon modifying the way this redox-energy state is expressed.

Because the metabolomics data pointed to remodeling at the interface between mitochondrial carbon metabolism and the urea cycle, we next examined whether IPA also affects CPS1-dependent urea cycle function. Glucagon modestly increased CPS1 activity, whereas IPA attenuated CPS1 activity most clearly under glucagon-stimulated conditions (Fig. 3D). IPA also reduced urea secretion in glucagon-stimulated hepatocytes (Fig. 3E), supporting a functional reduction in urea cycle output. Although *Cps1* mRNA expression was reduced in the IPA-treated cells (Fig. 3F), total CPS1 protein levels were not detectably decreased (Fig. 3D), indicating that the reduction in CPS1 activity is unlikely to be driven simply by reduced CPS1 abundance. Instead, these data suggest that IPA alters CPS1-dependent metabolism through changes in enzyme activity, substrate availability, or mitochondrial metabolic state.

Finally, pharmacological inhibition of CPS1 with H3B-120 reduced glucagon-stimulated glucose production, indicating that CPS1-dependent mitochondrial nitrogen metabolism contributes to the metabolic capacity required for maximal glucose output (Fig. 3G). However, IPA produced a further modest reduction in glucose production even in the presence of H3B-120. Thus, while CPS1 inhibition partially phenocopies the effect of IPA, the residual IPA sensitivity under CPS1-inhibited conditions suggests that IPA does not act solely through CPS1. Rather, CPS1-dependent urea cycle activity appears to represent one component of a broader IPA-sensitive mitochondrial metabolic state that supports lactate-dependent glucose production.

Together, these data indicate that IPA remodels the glucagon-stimulated mitochondrial state by altering Complex I-dependent redox balance, ATP availability, and CPS1-linked nitrogen metabolism, while limiting lactate-dependent hepatic glucose production through mechanisms that are only partly explained by reduced CPS1 activity.

### Endogenous IPA levels vary with nutritional state, and short-term IPA administration improves glucose handling in WD-fed mice

Given that IPA is a microbiota-derived metabolite generated from dietary tryptophan, we first examined whether systemic IPA availability varies across physiological nutritional states. In chow-fed mice with an intact microbiome (i.e., without antibiotic depletion), targeted LC-MS analysis revealed that circulating and hepatic IPA levels fluctuate with feeding status. Both serum and liver IPA concentrations differed between ad libitum, overnight fasted (16 h), and refed (2 h) conditions (Fig. 4A), consistent with the idea that endogenous IPA availability dynamically reflects nutrient intake and microbial metabolism of dietary tryptophan.

**Figure 4.**
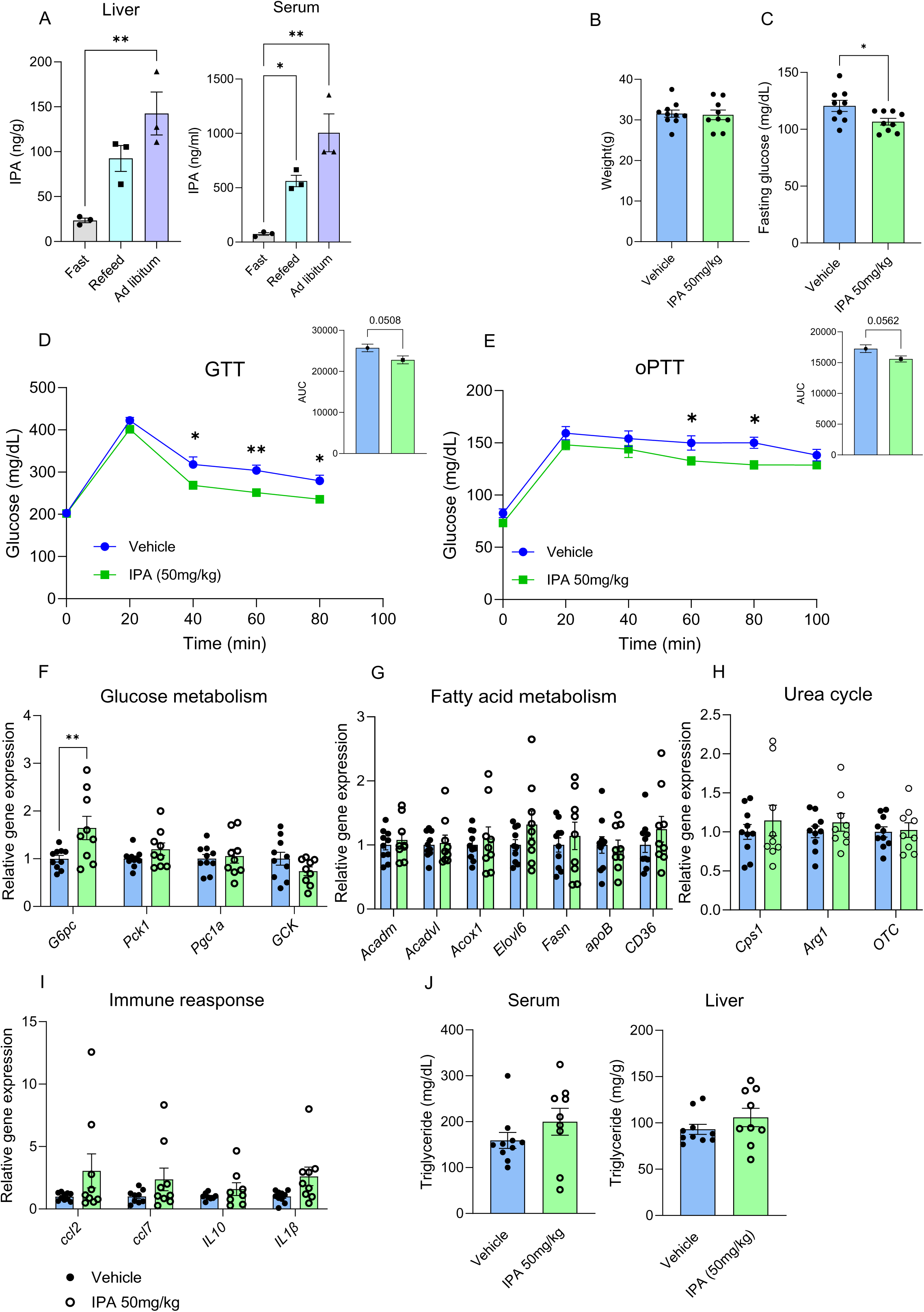
Acute IPA treatment improves metabolic homeostasis in vivo. **(A)** IPA concentrations in serum and liver under fasting, refeeding and ad libitum conditions. **(B)** body weight. **(C)** fasting blood glucose levels. **(D-E)** glucose tolerance test (GTT) and oral pyruvate tolerance test (oPTT). **(F-I)** hepatic gene expression analysis. **(J)** triglyceride levels in serum and liver. *, p<0.05; **, p<0.01.

Having established that IPA alters hepatocyte metabolic state and suppresses glucose production in vitro, we next asked whether increasing systemic IPA exposure influences glucose metabolism in vivo. Mice maintained on a Western diet (WD) for ∼8 weeks, also with an intact microbiome, were treated with IPA (50 mg/kg, i.p.) once daily for 5 days. Body weight remained unchanged during the treatment period (Fig. 4B), indicating no overt effects on energy balance. In contrast, fasting blood glucose was reduced in IPA-treated mice compared with vehicle-treated controls (Fig. 4C). During glucose tolerance testing, IPA-treated mice displayed reduced glycemic excursion, resulting in a lower GTT AUC (Fig. 4D). During pyruvate tolerance testing, IPA-treated mice also showed a reduced glucose excursion, reflected by a lower PTT AUC (Fig. 4E). These findings indicate that short-term IPA administration improves glucose handling in WD-fed mice and is associated with reduced pyruvate-driven glycemic excursion in vivo.

We next examined whether these physiological effects were accompanied by coordinated transcriptional remodeling of hepatic metabolic pathways. IPA treatment did not produce broad suppression of the hepatic gluconeogenic program (Fig. 4F). Instead, expression of several glucose metabolism genes was largely unchanged, while *G6pc* expression was increased (Fig. 4F). Genes involved in fatty acid metabolism were also mostly unchanged (Fig. 4G), indicating that short-term IPA exposure does not broadly reprogram hepatic lipid metabolic gene expression under these conditions. Similarly, urea-cycle and inflammatory genes (Fig. 4H, I) did not show a coordinated transcriptional pattern consistent with pathway-wide induction or suppression. Serum and hepatic triglyceride levels were not markedly altered (Fig. 4J), further suggesting that the short-term metabolic effects of IPA were not secondary to major changes in hepatic lipid accumulation or systemic triglyceride availability. Notably, this in vivo transcriptional profile differs from the suppression of glucagon-responsive gluconeogenic gene expression observed in primary hepatocytes treated with IPA (Fig 1). Thus, repression of the gluconeogenic transcriptional program is unlikely to represent the dominant mechanism underlying the in vivo phenotype. Instead, the reduction in fasting glycemia, improved glucose tolerance, and reduced PTT AUC are more consistent with changes in hepatic metabolic state or flux regulation rather than broad transcriptional reprogramming. Together with the in vitro data, these findings support a model in which IPA influences systemic glucose homeostasis, at least in part, by modulating hepatic metabolic function rather than by globally suppressing metabolic or inflammatory gene-expression programs.

### Microbial IPA production modestly improves glucose tolerance of chow-fed mice following microbiome depletion

To test whether microbial IPA production is sufficient to influence host glucose metabolism in vivo, we used a microbiome-depletion and recolonization approach based on *Clostridium sporogenes*, a gut bacterium previously shown to generate IPA through the *fldC*-dependent reductive metabolism of tryptophan^30^. Antibiotic-treated chow-fed mice were colonized with either wild-type *C. sporogenes* or an isogenic *fldC*-deficient strain that is impaired in IPA production, and glucose homeostasis was assessed following colonization (Fig. 5A). Consistent with the expected metabolic difference between the two strains, mice colonized with wild-type *C. sporogenes* exhibited higher circulating IPA levels than mice colonized with the *fldC*-deficient strain (Fig. 5B), while body weight was comparable between groups (Fig. 5C). Functionally, wild-type *C. sporogenes* colonization produced a modest but detectable improvement in glucose tolerance, reflected by lower glycemic excursion during GTT and a reduced GTT AUC compared with mice receiving the *fldC*-deficient strain (Fig. 5D). In contrast, pyruvate tolerance was largely unchanged between groups, with no clear reduction in PTT AUC (Fig. 5E). Thus, under chow-fed conditions, microbial IPA production is sufficient to modestly improve glucose handling, but does not strongly suppress pyruvate-driven glucose excursion in vivo.

**Figure 5.**
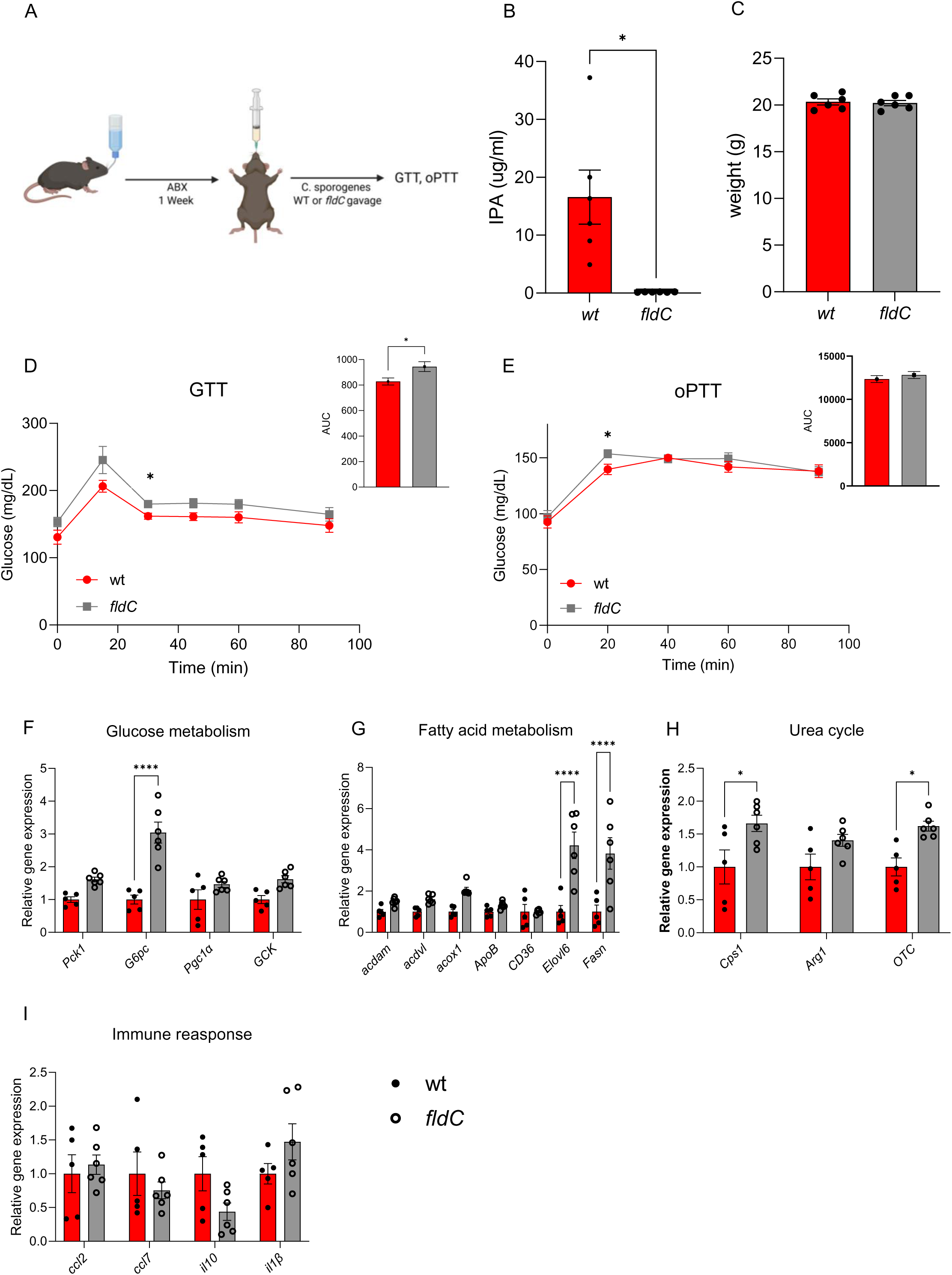
Microbial IPA production regulates metabolic phenotypes. **(A)** Experimental scheme of microbial recolonization. **(B)** Serum IPA levels following recolonization with WT or *fldC* bacteria lacking the capacity to synthesize IPA. **(C)** Body weight of recolonized mice. **(D-E)** Glucose tolerance test (GTT) and oral pyruvate tolerance test (oPTT). **(F-I)** Hepatic gene expression analysis. *, p<0.05.

We next examined hepatic gene expression in ad libitum-fed mice collected at the end of the recolonization experiment. This sampling condition was selected because endogenous microbial IPA production is expected to depend on dietary substrate availability and may therefore be more apparent in the fed state. Under these conditions, colonization with wild-type *C. sporogenes* altered the expression of selected genes involved in glucose metabolism (Fig. 5F), fatty acid metabolism (Fig. 5G), urea cycle function (Fig. 5H), and immune response (Fig. 5I), but the overall pattern did not indicate a uniform suppression of the hepatic gluconeogenic program. Importantly, these samples differ from the acute IPA administration experiment in WD-fed mice, in which animals were fasted overnight, injected with IPA the following day, and sacrificed two hours after the final injection. Thus, differences between the transcriptional profiles in the two in vivo settings likely reflect both metabolic state and mode of IPA exposure: endogenous microbial production during ad libitum feeding versus acute pharmacological IPA exposure after fasting. Consistent with this interpretation, the absence of a strong PTT phenotype in chow-fed recolonized mice suggests that microbial IPA production does not robustly suppress pyruvate-driven hepatic glucose output under these benign conditions.

Together with the direct IPA administration studies in WD-fed mice, these data provide complementary evidence that IPA can function as a microbiota-derived metabolic effector. The recolonization experiment strengthens the physiological relevance of IPA by linking *fldC*-dependent microbial production of the metabolite to improved glucose tolerance. However, the modest GTT effect and absence of a robust PTT phenotype indicate that, in chow-fed recolonized mice, microbial IPA production does not strongly suppress pyruvate-driven glucose excursion. This may reflect the relatively limited metabolic stress of the chow-fed state, the ad libitum-fed sampling condition, or the lower and more gradual exposure profile achieved through microbial production compared with acute IPA administration. The impact of microbial IPA production on hepatic glucose output may therefore become more evident under WD-induced metabolic challenge. Thus, the present data support a role for microbial IPA in systemic glucose homeostasis, while suggesting that its effects on hepatic glucose production depend on nutritional state, metabolic context, and mode of exposure.

## Materials and Methods

### Reagent

IPA (sigma-Aldrich, 220027), Indole (sigma-Aldrich, I3408), ICA (sigma-Aldrich, 284734), IAA (sigma-Aldrich, I5148), tryptamine (chemcruz, sc-206065), IBA (sigma-Aldrich, I5386), Dexamethasone (Sigma, D4902), Glucagon (Sigma, G2044), insulin (Sigma, I-2643), sodium pyruvate solution 100 mM (Biological Industries, 03-041-1B). Sodium l-lactate (Sigma, L7022-10G). l-glutamine solution (Biological Industries, 03-020-1B). DMEM high glucose (Biological Industries, 01-052-1 A). DMEM low glucose (Biological Industries, 01-050-1 A). DMEM with no glucose, no glutamine, and no sodium pyruvate (Biological Industries, 01-057-1 A). Glucose (GO) assay kit (Sigma, GAGO20-1KT). Liberase Research Grade (Sigma, 5401119001). Lipofectamine RNAiMAX Transfection Reagent (Thermo Scientific, 13778075). Fetal bovine serum (Biological Industries, 04-007-1A). Lipofectamine RNAiMAX Transfection Reagent (Thermo Scientific, 13778075).

### Animals

6-8 weeks old C57BL/6JOlaHsd males were purchased from Envigo, and housed under specific pathogen-free (SPF) conditions. The mice were kept in environments with 40–70% relative humidity, ambient temperature of 22(± 2)°C and a 12 h/12 h light/dark period. Where indicated, mice were fed high fat/high sucrose diet (Tekald, TD.08811). All animal procedures were approved by the Hebrew University of Jerusalem IACUC. IPA was dissolve In DMSO to concentration of 100mg/ml. the working solution was prepared by mixing 10% IPA stock or DMSO, 10% Tween 80 and 80% PBS. The solution was administered at 50 mg/kg body weight.

### Glucose and pyruvate tolerance tests

Mice were fasted overnight. For PTT, mice were orally gavaged with a 25% pyruvate solution in Dulbecco PBS (dPBS) at a dose of 2.5 g/kg. For GTT, mice were gavaged with a 20% D-glucose solution in dPBS at a dose of 2 g/kg. Blood glucose levels were measured at the indicated time points following injection.

### Antibiotic administration

1g/L ampicillin(Formedium), 1g/L Metronidazole(Sigma-aldrich, M3761), 1g/L Neomycin sulfate(goldbio 1405-10-3), 0.5g/L vancomycin (ApexBio B1223) were administered in the drinking water for a week.

### *C. sporogenes* colonization

*Clostridium sporogenes* wild-type (WT) and *fldC* mutant strains were kindly provided by the Sonnenburg lab^30^. Bacteria were cultured in an anaerobic chamber (COY labs, USA) overnight in TYG medium (30 g/L tryptone, 20 g/L yeast extract, 1 g/L sodium thioglycolate). Cultures were supplemented with glycerol to a final concentration of 30% and stored at −80°C. Following antibiotic treatment, mice were orally gavaged with approximately 8 × 10□ bacteria daily for one week. To assess bacterial abundance, fresh stool samples from each cage were inoculated into TYG medium and incubated overnight under anaerobic conditions. The following day, bacterial cultures were centrifuged at 10,000 × g for 10 minutes, and the resulting pellets were resuspended in 500 μL DDW. Samples were then heated at 95°C for 10 minutes to lyse the bacteria and release DNA. PCR amplification was performed using specific primers and the Phanta DNA polymerase enzyme, according to the manufacturer’s instructions. PCR products were analyzed by electrophoresis on a 1% agarose gel.

### Primary mouse hepatocyte isolation and culture conditions

Primary hepatocytes were isolated using a Liberase perfusion technique. Eight-week-old WT mice were sacrificed by Isoflurane inhalation, and the liver was perfused via the vena cava with HBSS (0.5 mM EDTA, 25 mM HEPES, calcium-free, magnesium-free, phenol red-free; Sartorius Israel, 02-018-1A). During perfusion, the portal vein was clamped for 5 seconds after being cut. Subsequently, 12.5 mL of digestion buffer containing 25mg/ml Liberase was perfused. The liver was excised, rinsed with 10 mL of dPBS (Sartorius Israel, 02-023-1A), and gently scraped. The tissue was suspended in 20 mL of plating medium and passed through a 70- μm cell strainer. Live hepatocytes were separated using a Percoll gradient and plated in plating medium (high-glucose DMEM supplemented with 10% FBS, 2 mM pyruvate, 1% penicillin/streptomycin, 0.1 μM insulin, and 1 μM dexamethasone). After 3-4 hours the medium was replaced with maintenance medium (low-glucose DMEM supplemented with 0.2% BSA, 4 mM L-glutamine, 1 mM pyruvate, 1% penicillin/streptomycin, 1 nM insulin, and 0.1 μM dexamethasone). For siRNA-mediated suppression of PXR, cells were transfected with siRNA oligos targeting *Nr1i2* (purchased for IDT) or a non-targeting control according to the manufacturer’s instructions. After 48 hours of incubation, knockdown efficiency was assessed by quantitative PCR.

### Hepatic Glucose Production

Day after isolation, primary hepatocytes were incubated for 3 hours with starvation medium (low-glucose DMEM supplemented with 0.2% BSA, 4 mM L-glutamine, 1 mM pyruvate, 1% penicillin/streptomycin) containing treatment. Following that the cells washed with dPBS and the media was replaced to DMEM devoid of glucose and phenol red (Sigma D-5030), supplemented with 0.2% BSA, sodium bicarbonate, and 1% Penicillin/Streptomycin. Additionally, the medium contained either sodium pyruvate and sodium L-lactate(2mM and 20mM, respectively) Glycerol (10mM) or Glutamine (4mM) as gluconeogenic substrates. Cells were exposed to these conditions for 6 hours with glucagon (100nM) stimulation. For the glucose quantification, 50 μl of the cell culture media was collected and mixed with 100 μl of the Glucose Oxidase (GO) Assay Kit (Sigma, GAGO20-1KT). The samples were then incubated at 37°C for 30 minutes. To terminate the reaction, 100 μl of 12N sulfuric acid was added. The absorbance of the resulting glucose was measured at 540 nm. Glucose levels were normalized to total protein.

### Gene expression

In the morning, culture medium was replaced with hormonal starvation medium containing the indicated treatment for 3 h. Cells were then stimulated with glucagon (100 nM) for an additional 3 h before harvesting.

### RNA Extraction and cDNA preparation

Total RNA was isolated using one of two methods. Cell culture total RNA was isolated using Nucleospin RNA, mini kit for RNA purification (Macherey-Nagel, #740955.50), according to the manufacturer’s protocol. Total RNA from liver tissue was extracted using TRIzol (or TRI reagent). Subsequently, cDNA was synthesized using a qScript cDNA Synthesis Kit (Quantabio, #95047-500) following the manufacturer’s instructions.

### Quantitative real-time Polymerase Chain Reaction (qPCR)

qPCR was performed with a C1000 Touch thermal cycler CFX384 instrument (Bio-Rad) using Luna Universal qPCR master mix (NEB, M3003E). All qPCR experiments were replicated at least three independent times. Gene values were normalized with *b2M* and *Rpl13*. Bio-Rad CFX Maestro software was used to analyse gene expression.

### Metabolite extraction for polar metabolite analysis

Extraction and analysis of polar metabolites and lipids were conducted following previously established methods^31,32^, with minor modifications. Mouse primary hepatocytes (2×10^6^ cells), treated with indole-3-propionic acid (IPA) for 3 hrs or with DMSO (control), were extracted with 1 mL of pre-chilled (−20°C) MTBE:methanol (1:3, v/v). Samples were vortexed briefly, sonicated for 30 min in an ice-cold bath (with intermittent vortexing every 10 min), and then mixed with 0.5 mL of DDW:methanol (3:1, v/v) containing labeled amino acid standards (^13^C and ^15^N; Sigma, 767964; 1:500 dilution). After thorough vortexing and centrifugation, the upper organic phase was collected. The remaining polar phase was extracted using an additional 0.5 mL MTBE. Polar phase extracts were collected and dried under a gentle nitrogen stream, and stored at −80°C.

### LC-MS Analysis of Polar Metabolites

Dried polar extracts were reconstituted in 120 µL methanol:DDW (50:50, v/v), centrifuged twice, and 70 µL was transferred to UPLC vials. Analysis utilized an Acquity I-class UPLC system (Waters) coupled with a Q Exactive Plus Orbitrap mass spectrometer (Thermo Fisher Scientific) operating in negative ionization mode. Metabolites were separated using a SeQuant Zic-pHilic column (150 × 2.1 mm) with a corresponding guard column (Merck). Mobile phases comprised ACN (phase B) and 20 mM ammonium carbonate with 0.1% ammonium hydroxide in DDW:ACN (80:20, v/v; phase A). The flow rate was 200 µL/min at 45°C. The gradient was: 75% B (0–2 min), decreasing to 25% B by 14 min, holding until 18 min, returning to 75% B at 19 min, and equilibration until 23 min. Injection volume was 2 µL. Mass spectra were acquired in negative heated electrospray ionization (HESI) mode over the mass range 70–1050 m/z. Source parameters: capillary temperature 325°C, spray voltage 3.25 kV, sheath gas 40 units, auxiliary gas 10 units, auxiliary gas temperature 50°C. MS1 spectra were acquired at a 35,000 FWHM resolution, and MS/MS data collection was performed at a 17,500 FWHM resolution (3 m/z isolation window). Polar data processing was done using Progenesis QI (Waters). identifying compounds based on accurate mass (≤5 ppm), retention time, and spectral libraries generated from injected standards. Relative polar metabolite levels were normalized to the internal standards, and cell number.

### ^13^C-lactate tracing

Primary mouse hepatocytes were treated with IPA (50μM) overnight and then hormonally starved for 3 h, still in the presence of IPA. Then, they were incubated for 3 h in glucose-production medium containing [U-^13^C]-lactate (Cambridge Isotopes), with vehicle or IPA in the absence or presence of glucagon. Cells were washed with ice-cold PBS, and metabolites were extracted under cold conditions using 80% Methanol solution. Polar metabolites were analyzed by ion chromatography–high-resolution mass spectrometry, using a workflow adapted for polar organic acids and phosphorylated intermediates. Lactate-derived carbon incorporation was determined from isotopologue enrichment in glucose-6-phosphate and TCA-cycle intermediates, with correction for natural isotope abundance.

### Targeted measurement of IPA in serum and liver

IPA levels were measured in serum and liver samples by targeted LC-MS/MS. Serum samples and homogenized liver extracts were processed using LC-MS-compatible extraction conditions, cleared by centrifugation, and analyzed using a UPLC system coupled to a triple-quadrupole mass spectrometer. Chromatographic separation was performed on a C18 column using an acetonitrile/water gradient, and IPA was detected by electrospray ionization in scheduled MRM mode, using an analytical workflow adapted from the LC-MS/MS method for plant hormones and growth regulators. Data were analyzed using targeted quantification software and normalized to sample volume or tissue weight, as appropriate.

### Western blot

Cells were incubated in hormonal starvation medium containing the indicated treatments for 3 hours. Following this, the medium was replaced with fresh hormonal starvation medium containing the same treatments and either glucagon (100 nM) or insulin (20 nM) for 30 minutes. After treatment, cells were harvested in RIPA buffer supplemented with protease inhibitors and shaken at 4°C for 15 minutes. Cells were then scraped and transferred to Eppendorf tubes, followed by centrifugation at maximum speed at 4°C for 15 minutes. The supernatant was collected into new Eppendorf tubes. Protein concentration was determined, and 18 µg of protein from each sample was separated by 10% SDS-PAGE and transferred to PVDF membranes (Millipore IPVH00010). Membranes were blocked and incubated overnight at 4°C with the following primary antibodies: CREB (CST, 9104S), p-CREB (CST, 9198S), AKT (CST, 2920S), p-AKT (CST, 4060S), AMPK (CST, 5831), p-AMPK (CST, 2535), and Vinculin (CST, 13901). After washing with TBST, membranes were incubated with peroxidase-conjugated secondary antibodies in 5% skim milk at room temperature for 1 hour. Protein signals were detected using enhanced chemiluminescence substrate (ECL, Bio-Rad).

### Intracellular NAD^+^/NADH

Intracellular NAD□ and NADH levels were measured using an enzymatic cycling assay as previously described^33^. Briefly, 2.5 × 10□ cells were seeded in 10-cm culture dishes. Cells were treated with IPA for 3 hours, followed by stimulation with glucagon for an additional 3 hours. During the final 1 hour of incubation, either rotenone (25 nM) or oligomycin (2 μM) was added to the culture medium. Following treatment, cells were washed with cold dPBS, scraped into 5 ml dPBS, and centrifuged at 1,000 × g for 5 minutes at 4°C. The cell pellet was resuspended in 500 μl extraction buffer (20 mM nicotinamide, 20 mM NaHCO□, 100 mM Na□CO□) and centrifuged at 1,200 × g for 15 minutes at 4°C. For NADH determination, 250 μl of the supernatant was incubated at 60°C for 30 minutes to decompose NAD□. Both heated (NADH) and unheated (total NAD) samples were used for analysis. A reaction mixture containing 160 μl NAD-cycling buffer (100 mM Tris-HCl, pH 8.0; 0.5 mM thiazolyl blue; 1 mM phenazine ethosulfate; 1% BSA) and 5 unit alcohol dehydrogenase (ADH) was added to each well of a 96-well plate containing 20 μl of sample. After incubation for 1 minute at 30°C in the dark, absorbance at 570 nm was measured. Subsequently, 20 μl of 6 M ethanol was added to initiate the cycling reaction, and the change in absorbance at 570 nm was recorded every 30 seconds for 10 minutes at 30°C using a plate reader. All samples were analyzed in triplicate. NAD□ levels were calculated by subtracting NADH values from total NAD measurements.

### Mitochondrial isolation and CPS1 activity

A total of 2.5 × 10□ cells were seeded in 10-cm culture dishes. Cells were treated with IPA for 3 hours, followed by stimulation with glucagon for an additional 3 hours. Following treatment, cells were washed with ice-cold dPBS, scraped into 5 mL dPBS, and centrifuged at 1,000 × g for 5 minutes at 4°C. The resulting cell pellet was resuspended in 1 mL ice-cold hypotonic Tris buffer (10 mM, pH 7.6) and transferred to 2 mL microcentrifuge tubes.

After a 3-minute incubation on ice, cells were homogenized using a homogenizer (1,000 rpm, 15 strokes). Subsequently, 200 μL of 1.5 M sucrose was added to the homogenate, followed by centrifugation at 600 × g for 10 minutes at 4°C. The supernatant was collected and transferred to a new tube, then centrifuged at 14,000 × g for 10 minutes at 4°C to pellet mitochondria. The supernatant was discarded, and the mitochondrial pellet was resuspended in 200 μL of 10 mM Tris buffer (pH 7.6). For CPS1 activity measurement^34^, 5 μg of mitochondrial protein was incubated in a final volume of 100 μL containing assay buffer (50 mM HEPES, pH 7.0; 12.5 mM KHCO□; 0.75 mM (NH□)□SO□; 5 mM MgCl□; 50 mM KCl; 1 mM TCEP; 0.01% Triton X-100; 0.0025% BSA; 0.2 mM N-acetylglutamate; 5 mM ATP). Samples were incubated at 37°C for 1 hour. Following incubation, 40 μL of the reaction mixture was transferred to a PCR tube, and 2 μL of 2 M hydroxylamine was added. Samples were heated at 95°C for 10 minutes. Equal volumes of solution A (0.85 g antipyrine dissolved in 40% H□SO□) and solution B (0.625 g diacetyl monoxime dissolved in 5% acetic acid) were mixed, and 160 μL of this mixture was added to each sample. The samples were then incubated at 95°C for 15 minutes. After cooling to room temperature, absorbance was measured at 458 nm using a plate reader.

### Urea detection

A day after seeding, cells were treated with IPA or DMSO for 3 hours under hormonal starvation conditions. Following starvation, cells were washed twice with DPBS and incubated for 6 hours in phenol red–free DMEM-based assay medium supplemented with 8.3 mg/mL DMEM powder, 3.7 mg/mL sodium bicarbonate, 0.2% BSA, 5 mM glucose, 1% penicillin–streptomycin, 2 mM sodium pyruvate, 2 mM L-glutamine, and 1 mM NH□Cl, in the presence or absence of IPA and glucagon. Urea concentration was measured in cell culture supernatants, which were collected after the incubation period, and in serum samples, using a commercially available assay kit (abcam, ab234052) according to the manufacturer’s instructions.

### Triglyceride detection

Approximately 50 mg of liver tissue was homogenized in 5% NP-40 using bead beating. Samples were then heated to 100°C and cooled to room temperature twice, followed by centrifugation at maximum speed for 2 minutes. The resulting supernatants were collected and diluted 1:10 prior to analysis. Serum samples were diluted 1:5. Triglyceride concentration was measured using a commercially available assay kit (abcam, ab65336) according to the manufacturer’s instructions..

### ATP measurement

Cells were seeded in 24-well plates and, the following day, the medium was replaced with hormonal starvation medium containing IPA or DMSO for 3 hours. After starvation, the medium was replaced with fresh medium containing the indicated treatments (IPA and glucagon). ATP levels were measured using a commercially available assay kit (Promega, G9241) according to the manufacturer’s instructions.

## Discussion

In this study, we identify indole-3-propionic acid (IPA) as a microbiota-derived indole metabolite that directly modulates hepatic glucose metabolism. IPA suppressed glucagon-stimulated glucose production in primary hepatocytes in a substrate-selective manner, with a stronger effect on pyruvate/lactate- and glutamine-supported glucose production than on glycerol-supported glucose production. This effect occurred without detectable inhibition of proximal glucagon-CREB or insulin-AKT signaling and was not prevented by perturbation of the canonical IPA-associated receptors AhR or PXR/*Nr1i2*. Mechanistically, IPA reduced the incorporation of lactate-derived carbon into gluconeogenic and mitochondrial intermediates, altered cellular NAD□/NADH redox balance through a Complex I-dependent process, increased ATP availability, and attenuated CPS1-linked urea cycle activity during glucagon stimulation. In vivo, short-term IPA administration improved glucose handling in WD-fed mice, while *fldC*-dependent microbial IPA production modestly improved glucose tolerance following microbiome depletion and recolonization. Together, these findings support a model in which IPA acts as a gut microbiota-derived metabolic effector that tunes hepatocyte mitochondrial metabolic state and thereby limits hepatic glucose output.

A central feature of the IPA response was its substrate selectivity. IPA robustly suppressed glucose production from pyruvate/lactate, while glycerol-supported glucose production was largely preserved. Because glycerol enters the gluconeogenic pathway downstream of mitochondrial pyruvate metabolism, this pattern argues against a nonspecific inhibition of gluconeogenesis or a general impairment of hepatocyte glucose-producing capacity. The additional suppression of glutamine-supported glucose production further supports the idea that IPA preferentially affects substrates that require mitochondrial metabolism, anaplerotic processing, or mitochondrial-cytosolic exchange before contributing to glucose synthesis. Although IPA also attenuated glucagon-induced *Pck1* and *G6pc* expression in primary hepatocytes, this transcriptional effect alone does not fully explain the functional phenotype. Glycerol-supported glucose production remained largely intact despite reduced expression of these genes, and the in vivo transcriptional profile did not show coordinated suppression of the gluconeogenic program. Thus, while IPA can influence selected glucagon-responsive transcripts, the dominant functional effect appears to involve metabolic control of substrate handling rather than broad transcriptional repression of gluconeogenesis.

The isotope tracing and metabolomics data provide further insight into this metabolic control. IPA reduced the abundance of ^13^C□-labeled G6P from [U-^13^C]-lactate, particularly under glucagon-stimulated conditions, directly linking the reduction in glucose output to diminished incorporation of lactate-derived carbon into the gluconeogenic pathway. IPA also reduced ^13^C□ enrichment in multiple TCA cycle intermediates, including citrate, α-ketoglutarate, fumarate, and malate. The coordinated nature of this reduction argues against inhibition at a single TCA cycle step and instead suggests altered entry, retention, or propagation of lactate-derived carbon within mitochondrial metabolism. Notably, these tracing results occurred alongside increased steady-state abundance of selected TCA-and urea cycle-associated metabolites, including citrulline, argininosuccinate, fumarate, and malate. This combination, reduced lactate-derived labeling but accumulation of selected metabolite pools, suggests that IPA does not simply suppress mitochondrial metabolism globally. Rather, IPA appears to remodel the source, turnover, or partitioning of metabolite pools at the interface between the TCA cycle, gluconeogenesis, and nitrogen metabolism.

The redox data suggest one mechanism by which IPA may alter lactate-dependent glucose production. IPA increased the cellular NAD□/NADH redox balance under basal conditions, and this effect was abolished by rotenone, indicating dependence on Complex I-linked mitochondrial electron transport. Glucagon also increased NAD□/NADH redox balance, consistent with activation of oxidative metabolism during gluconeogenic stimulation. However, IPA did not simply enhance this glucagon response. Instead, the combination of IPA and glucagon shifted the NAD□/NADH ratio toward baseline, indicating that IPA changes the characteristic redox response to glucagon. Since lactate-supported gluconeogenesis requires coordinated cytosolic and mitochondrial redox handling, altered NADH oxidation or redox coupling could limit efficient conversion of lactate-derived carbon into glucose even when upstream glucagon signaling remains intact. The increase in ATP levels, particularly during glucagon stimulation, further argues that IPA does not inhibit mitochondrial energy metabolism globally. Rather, IPA appears to induce a distinct mitochondrial metabolic state in which respiration-linked redox handling and energy availability are altered, but the routing of lactate-derived carbon into gluconeogenesis is reduced.

The CPS1 and urea-cycle findings extend this model beyond carbon metabolism alone. Untargeted metabolomics pointed to selective remodeling of metabolites linked to nitrogen handling, and IPA attenuated glucagon-stimulated CPS1 activity and urea secretion without a detectable reduction in total CPS1 protein. These data suggest that the reduction in CPS1-dependent metabolism is unlikely to reflect loss of enzyme abundance and may instead result from altered mitochondrial metabolic state, substrate availability, or post-translational regulation of CPS1 activity. Pharmacological inhibition of CPS1 reduced glucagon-stimulated glucose production and largely phenocopied the effect of IPA, while the combination of CPS1 inhibition and IPA did not produce a clearly additive suppression. This suggests that IPA-sensitive glucose production and CPS1-dependent urea-cycle activity converge on overlapping mitochondrial metabolic processes. Importantly, these findings do not necessarily imply that CPS1 is the direct molecular target of IPA. Rather, CPS1 activity may represent a functional node within the broader glucagon-stimulated mitochondrial program that supports lactate-dependent hepatic glucose production.

The in vivo experiments support the physiological relevance of this mechanism while also highlighting the rapid nature and context dependence of the IPA response. Previous studies have shown that IPA can improve metabolic^17,19^, inflammatory^21^, intestinal^18,19^, neurological^23^, or cardiovascular outcomes^20^ in vivo, but these effects have generally been examined after repeated exposure over days to weeks. For example, IPA improved fasting glucose, fasting insulin, and HOMA-IR in rats after six weeks of dietary supplementation^17^, and other studies have used repeated IPA administration or dietary supplementation over approximately 10 days to several weeks. In contrast, our IPA treatment paradigm was deliberately short, consisting of once-daily IPA administration for only five days in WD-fed mice. Despite this limited exposure window, IPA reduced fasting glucose and improved glucose handling during both GTT and PTT. These effects occurred without marked changes in body weight, serum or hepatic triglycerides, or coordinated transcriptional remodeling of hepatic lipid, inflammatory, urea-cycle, or gluconeogenic gene programs. Indeed, the hepatic transcriptional response in vivo differed from that observed in primary hepatocytes, reinforcing the idea that transcriptional repression of gluconeogenesis is unlikely to be the dominant mechanism underlying the physiological phenotype. Instead, the rapid improvement in fasting glycemia and reduced PTT AUC are more consistent with acute modulation of hepatic metabolic state or flux regulation. The *C. sporogenes* recolonization experiment provides complementary evidence that microbial IPA production can influence host glucose metabolism, as mice colonized with wild-type *C. sporogenes* had higher circulating IPA levels and modestly improved glucose tolerance compared with mice colonized with the *fldC*-deficient strain. However, pyruvate tolerance was largely unchanged under these chow-fed conditions. This difference between acute IPA administration in WD-fed mice and microbial IPA production in chow-fed recolonized mice likely reflects differences in metabolic stress, nutritional state, timing of sampling, and mode or magnitude of IPA exposure. Thus, microbial IPA production appears sufficient to improve systemic glucose handling, whereas stronger effects on pyruvate-driven glucose excursion may require a metabolically challenged state or higher acute IPA exposure.

Based on these findings, we propose that IPA links gut microbial metabolism to hepatic glucose regulation by altering the mitochondrial metabolic state of hepatocytes. In this model, IPA promotes respiration-dependent redox remodeling, modifies the handling of lactate-derived carbon within mitochondrial and gluconeogenic pathways, and attenuates glucagon-stimulated CPS1-linked urea-cycle activity. These coordinated effects reduce the capacity of mitochondria-dependent substrates to support hepatic glucose production without globally inhibiting gluconeogenesis, disrupting proximal hormone signaling, or broadly suppressing hepatic metabolic gene expression. This model places IPA within a growing class of microbiota-derived metabolites that act not only through classical receptor-mediated pathways, but also by directly reshaping intracellular metabolic networks. From a therapeutic perspective, these findings suggest that microbial tryptophan metabolism, IPA-producing bacteria, or optimized IPA-related compounds could potentially be used to modulate excessive hepatic glucose output in metabolic disease. The relatively rapid improvement in glucose handling after short-term IPA administration further supports the possibility that this pathway may be pharmacologically or microbiologically actionable. However, several issues remain to be addressed before such approaches can be advanced, including the durability of the response, the exposure levels required to affect hepatic metabolism, the contribution of extrahepatic tissues, and whether sustained enhancement of IPA production can improve glucose homeostasis under diet-induced metabolic stress.

In summary, our findings identify IPA as a microbiota-derived regulator of hepatocyte mitochondrial metabolic state. By selectively limiting lactate- and glutamine-supported glucose production, remodeling redox and ATP balance, and attenuating CPS1-linked mitochondrial nitrogen metabolism, IPA directly modulates the metabolic programs that support glucagon-stimulated hepatic glucose output. These results provide a mechanistic framework for understanding how microbial tryptophan metabolites can influence systemic glucose homeostasis through direct effects on liver metabolism.

## Acknowledgments

This work was supported by EFSD/Boehringer Ingelheim European Research Programme on “Multi-System Challenges in Diabetes”, and by the HUJI-Hohenheim collaborative grant. We thank all members of the laboratory for their valuable discussions and technical assistance throughout the study.

---

**Figure S1.**
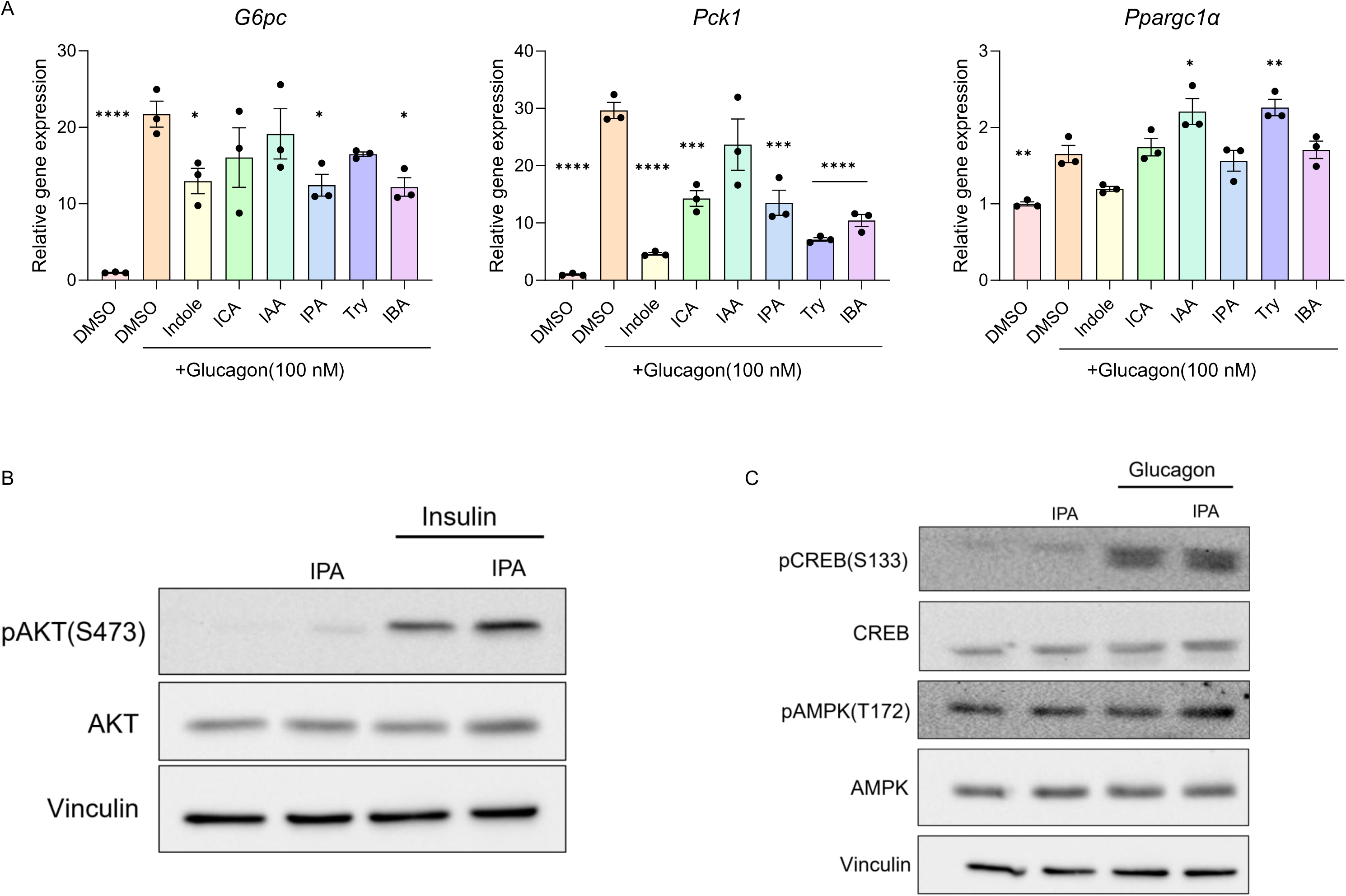
Indole-derived metabolites suppress gluconeogenic genes but do not modulate hepatic signaling. **(A)** Hepatic gene expression analysis of indole-derived metabolites and related targets. For statistics, the comparison was made to DMSO + glucagon conditions. **(B)** Western blot analysis of p-AKT under insulin stimulation. **(C)** Western blot analysis of p-CREB and p-AMPK under glucagon stimulation. *, p<0.05; **, p<0.01; ***, p<0.001; ****, p<0.0001.

**Fig S2.**
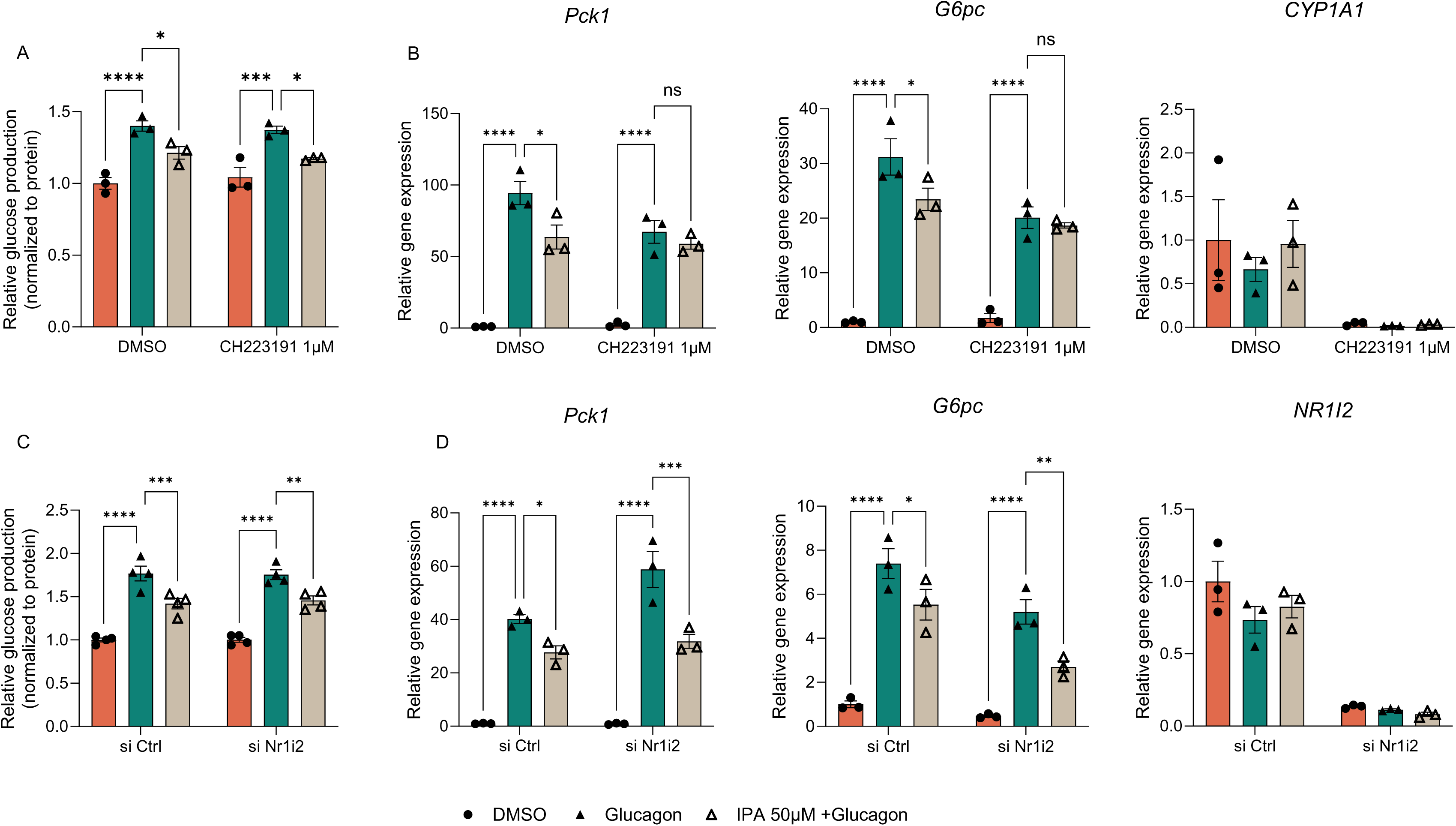
IPA effects persist Independent of AhR and PXR (*NR1I2*). (A-B) Hepatic glucose production and gene expression following AhR inhibition with CH223191. *Cyp1a1* is an AhR target gene. **(C-D)** Hepatic glucose production and gene expression following siRNA-mediated NR1I2 (encoding PXR) knockdown. *, p<0.05; **, p<0.01; ***, p<0.001; ****, p<0.0001.

